# Temporal, spatial, and parasitic drivers of microbial variation in European honey bees

**DOI:** 10.64898/2026.07.10.737668

**Authors:** Victor Rossier, Thibault Leroy, Philipp Engel, Markus Neuditschko, Vincent Dietemann, Benjamin Dainat

## Abstract

Although the roles of host-associated microbiomes in animal health are increasingly recognised, the factors influencing their variation remain understudied. The relatively simple microbiome of honey bees is a relevant system to address this gap. In particular, the relationship between variations in microbiome composition and the ectoparasite *Varroa destructor*, the main threat to honey bee health worldwide, is poorly established. In this study, we used metagenomic and statistical analyses of 1442 European honey bee colonies to investigate the relationships between the honey bee microbiome, temporality, location, *V. destructor* load, and behavioural response to its infestation by the host. While season, year, and location were identified as the main drivers of microbiome variation, *V. destructor* load emerged as a significant factor associated with microbiome variation. Notably, we identify several pathogens and opportunists that correlated positively with *V. destructor* load, while the core symbiont *Bombilactobacillus* correlated negatively. This is compatible with a shift in the microbiome toward dysbiosis, which may be driven by or promote *V. destructor* parasitism. By contrast, we found only limited evidence of an association between the microbiome and resistance behaviours of the host against this parasite. While the study cannot establish causal relationships, we present the largest metagenomic analysis of honey bee microbiomes to date, providing robust, generalisable evidence about the factors driving variation in the microbiome composition of this ecologically and economically important pollinator. These findings may serve as additional markers in selective breeding programs targeting *V. destructor* resistance, which could ultimately improve honey bee health.

## Introduction

Host-associated microbiomes are increasingly recognized as major determinants of animal health (Peixoto et al., 2021). Symbiotic microorganisms can enhance host resilience through multiple mechanisms, including food digestion, nutrient provisioning, immune modulation, and protection against pathogens (Lynch & Hsiao, 2019). Moreover, microbiome plasticity and host-microbiome interactions enable rapid responses and adaptation to changing environments (J. Li & King, 2025; Voolstra & Ziegler, 2020). Upon dysbiosis, the disruption of the balanced host-microbiome state, beneficial microbes often decline in favour of opportunistic pathogens, undermining host health and contributing to diseases (Caballero-Flores et al., 2023). Despite a growing interest in host-microbiome interactions, further research is needed to identify the sources of microbiome variation, particularly in case of dysbiosis, in order to improve animal health and resilience (Peixoto et al., 2021).

The microbiome of the honey bee *Apis mellifera* (Fig. 1) is a relevant and tractable system for studying how microbiome variation, including dysbiosis, influences host health (Zheng et al., 2018). Its relevance stems from the ecological importance of honey bees as essential pollinators (Klein et al., 2007), whose health is increasingly threatened (Goulson et al., 2015; Popovska Stojanov et al., 2021) and strongly influenced by their microbiome (Motta & Moran, 2024; Raymann & Moran, 2018). The honey bee microbiome is tractable because of the limited complexity of its gut microbiota, which is dominated by five core genera and involved in nutrient digestion and detoxification (Kešnerová et al., 2017; Zheng et al., 2016, 2019), modulation of immune system responses (Horak et al., 2020), mediation of pathogen defense (Steele et al., 2021), and enhancement of learning, memory (Zhang, Mu, Cao, et al., 2022), and social interactions among nestmates (Liberti et al., 2022; Zhang, Mu, Cao, et al., 2022). In addition to these gut symbionts, a wide diversity of pathogens has been described in and on honey bees (Engel et al., 2016), including bacteria (Forsgren, 2010; Genersch, 2010; Lang et al., 2022; Mouches et al., 1983; Raymann et al., 2018), fungi (Higes et al., 2007; Schaub, 1994; Aronstein & Murray, 2010), trypanosomid protists (Higes et al., 2007; Schaub, 1994), and viruses (Hartmann et al., 2015; Martin & Brettell, 2019). Under dysbiotic conditions, these microorganisms can proliferate and contribute to increased colony mortality (Maes et al., 2016). Yet, despite the honey bee microbiome being one of the best characterised host-associated microbial systems, large-scale studies examining the drivers of its variation and the consequences of this variation for host health remain scarce.

**Figure 1.**
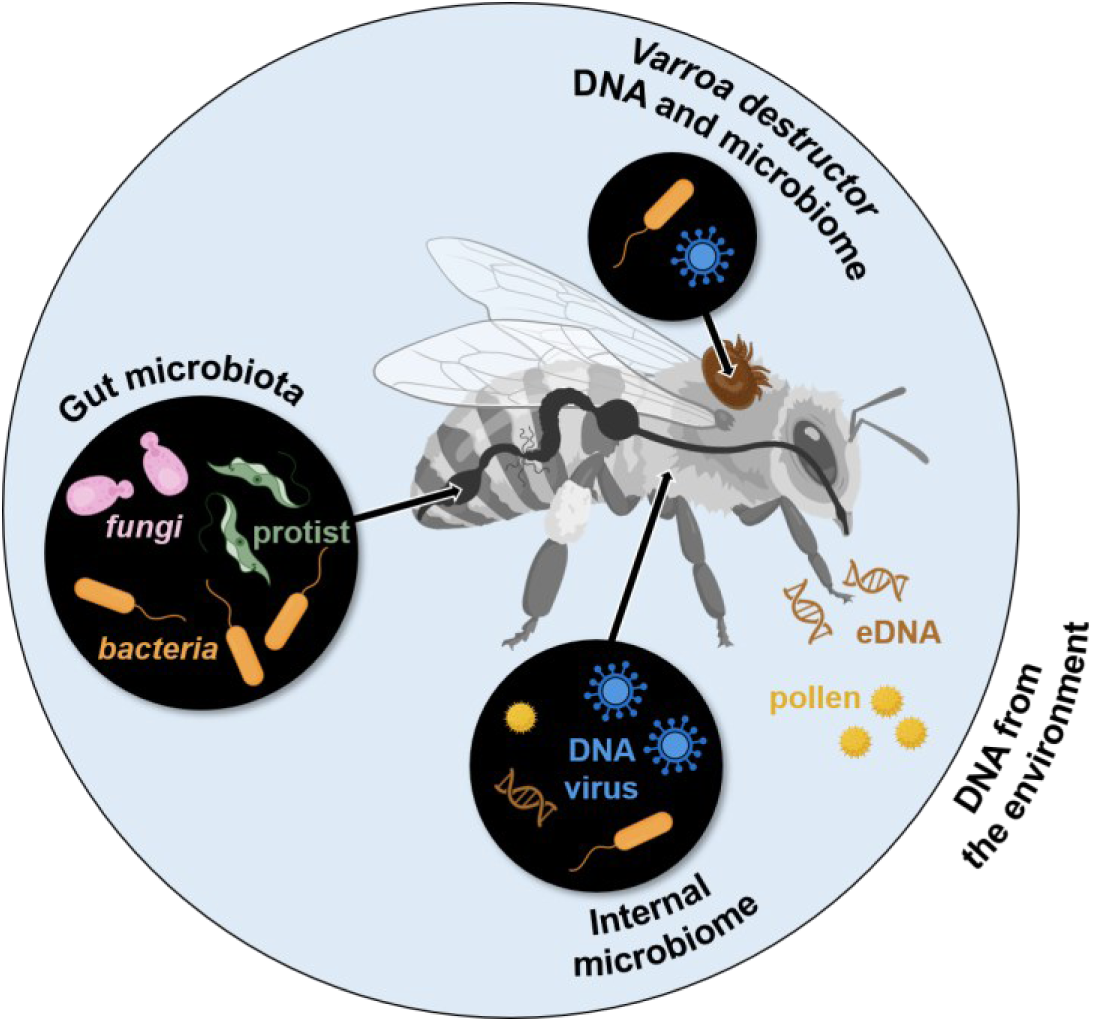
The honey bee-associated microbiome obtained from genomic data of pooled whole workers includes i) the gut microbiota and microorganisms found in the honey bee haemolymph such as DNA viruses and bacteria (internal microbiome); ii) DNA from the environmental (eDNA) such as pollen from plants or DNA from various organisms, which can be found on the surface of the honey bees and the mites or in their digestive tract; iii) the ectoparasitic mite *Varroa destructor*, including its microbiome. This broad microbiome definition is inspired from Berg et al., 2020. Figure created using BioRender.com.

Primarily through its role as a vector of viruses and bacterial pathogens, the ectoparasite mite *Varroa destructor* is the most serious threat to honey bees worldwide (Dietemann et al., 2012; Traynor et al., 2020; Morfin et al., 2023). The lifecycle of this parasite and its impact on honey bee health makes it a potential major factor of microbiome variation and dysbiosis. By feeding on honey bee haemolymph and fat body tissues (Han et al., 2024; Piou et al., 2023), *V. destructor* mites may facilitate the transmission of microorganisms (Hubert et al., 2015, 2016). Beyond the mere vectoring role of *V. destructor*, several studies have explored the links between this parasite’s load and the honey bee microbiome composition (Arredondo et al., 2025; Hubert et al., 2017; Kim et al., 2023; Y. Wang et al., 2025). Notably, lower *V. destructor* loads have been consistently associated with higher abundances of *Lactobacillidae* spp. (Hubert et al., 2017; Pitek et al., 2025; Tejerina et al., 2020; Y. Wang et al., 2025). By contrast, associations involving other taxa remain inconsistent across studies, both in direction and magnitude (Hubert et al., 2017; Y. Wang et al., 2025). These discrepancies likely reflect the limited sample sizes and restricted geographic and temporal coverage considered, which hinder the generalization of current findings about *V. destructor*-microbiome interactions (Hubert et al., 2016).

Microbiome variation may also be associated with the various resistance behaviours expressed by honey bee colonies against *V. destructor* (Guichard et al., 2020; Mondet et al., 2020; Svobodová et al., 2023). These include uncapping and recapping of brood cells (a trait called recapping), hygienic behaviour involving the removal of infected brood, and disruption of mite reproduction, which can be quantified directly via the decreased mite reproduction trait (DMR, von Virag et al., 2022). Although resistance traits are thought to involve neuronal, olfactory, developmental, and immune pathways, the limited overlap in associated genetic markers found across studies (Eynard et al., 2024; Guichard et al., 2022; Mondet et al., 2020) suggests other underlying mechanisms. The gut microbiota could play a role in the expression of these traits via the microbiota-gut-brain axis, whose implication in the regulation of insect behaviors has recently been revealed (Liberti et al., 2024; Liberti & Engel, 2020). Indeed, microbial metabolites produced in the gut can modulate the host neurophysiology by influencing immune, neuronal, or endocrine pathways beyond the gut. In honey bees, the gut microbiota has been shown to increase the proportion of neuroactive metabolites in the brain and affect pathways underlying sensory perception, which is essential for most honey bee behaviours (Cabirol et al., 2024; Liberti et al., 2022; Zhang, Mu, Shi, et al., 2022), including *V. destructor* resistance (Mondet et al. 2020; Mondet et al., 2021; Wagoner, 2023). Apart from three recent studies focusing on hygienic behaviour, the impact of *V. destructor* resistance traits on honey bee microbiome variation remains largely unexplored (De Iorio et al., 2026; Tola et al., 2025; Wang et al. 2026).

Here, we exploit the largest publicly available genomic dataset of pooled honey bee workers (Eynard et al., 2024) to identify factors (temporal, spatial, and linked to *V. destructor* parasitism) explaining honey bee microbiome variation. Since these samples consist of whole bees, they encompass both host and associated microbial DNA. Thus, by performing metagenomic taxonomic classification, we first provide a large-scale profiling of the honey bee microbiome across five European countries. We then quantify the contribution of season, year, location, and *V. destructor*-related phenotypes (*V. destructor* load, recapping and DMR) to microbiome variation. Although to a lesser extent than temporality and location, we identify *V. destructor* load as a key factor associated with variations of the honey bee microbiome composition. To understand which parts of the microbiome are associated with *V. destructor*-related phenotypes, we searched for associations between microbial taxa at the genus level and these phenotypes. We find diverse pathogenic and opportunistic genera associated with higher mite load and a core symbiont associated with lower mite load. This suggests a shift in the composition of the honeybee microbiome towards a dysbiotic state associated with parasitism by *V. destructor*. Overall, this large-scale metagenomic study provides robust and generalizable insights into the sources of honey bee microbiome variations highlighting microbial signatures linked to host vulnerability and resilience.

## Methods

In this study, we reanalysed the genomic data (300-500 pooled honey bee workers per colony) and *V. destructor*-related colony phenotypes (*V. destructor* load, recapping, and DMR) from 1513 colonies sampled in France, Switzerland, Netherlands, Luxembourg, Sweden, and New Zealand by Eynard et al. (2024; Fig. 2). Compared to the original study, our work explicitly focused on the non-bee sequencing reads, *i.e.* those that did not map to the honey bee genome in order to identify associations between the honey bee microbiome (Fig. 1), the *V. destructor*-related phenotypes, year, season, and location. Because preliminary analyses revealed the microbiomes of New Zealand colonies to be outliers, we restricted our analyses to the 1442 European colonies of the dataset. The final dataset comprised colonies from 149 apiaries sampled once between June and October in 2016, 2017 or 2018 (Supp. Table 1).

**Figure 2.**
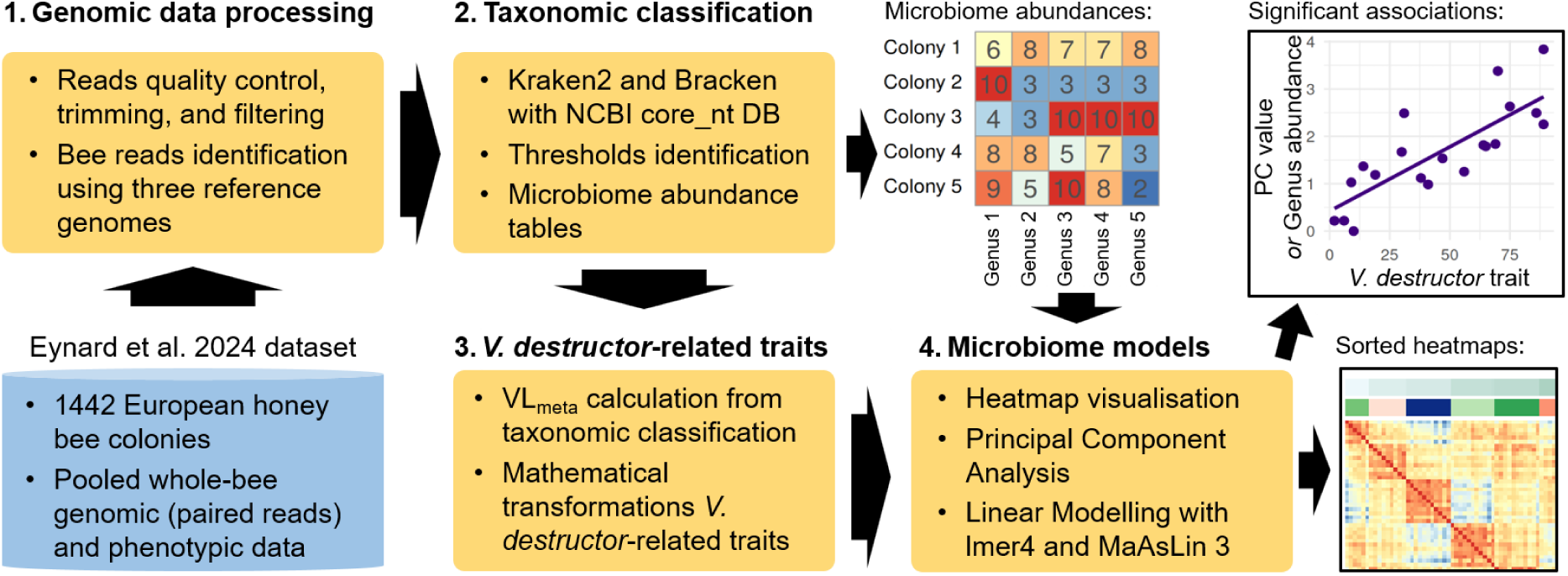
Overview of the bioinformatic pipeline used in this study. VL_count_: *V. destructor* load quantified by washing adult bees with detergent; VL_meta_: genomic-derived metric of *V. destructor* load; PC: principal component.

### Genomic data preprocessing

Raw DNA libraries were downloaded from NCBI and their quality was assessed using FastQC v0.12.1 and MultiQC v1.28. 3. Adapter sequences (TruSeq3-PE-2) and poly-G tails, defined as at least ten consecutive guanines and a known Illumina sequencing artifact (Andrew 2016), were removed using Cutadapt v4.4, and resulting reads shorter than 50 bp were then discarded.

Honey bee reads were identified by aligning libraries using Bowtie2 v2.5.1 against three reference honey bee genomes from diverse European lineages: the *A. mellifera* reference genome Amel_HAv3.1 (GCA_003254395.2; DH4 line with mixed European ancestry; Wallberg et al., 2019), *A. mellifera mellifera* (GCA_003314205.2), and *A. mellifera caucasica* (GCA_013841205.1, Eynard, Klopp, et al., 2024; Yokoi et al., 2022). Completely unmapped paired reads, *i.e.* for which neither mate mapped to any of these reference genomes, were assumed to be non-bee reads. They were extracted using Samtools v1.19.2 (*i.e.* samtools view -f 12) and used in the subsequent metagenomic analyses.

### Metagenomic taxonomic classification

Next, the non-bee reads were taxonomically classified using Kraken2 v.2.1.3, and abundance estimation was performed at the genus level using Bracken v2.9 (Fig. 2.1, Lu et al., 2017, 2022; Wood et al., 2019). Because of the taxonomic diversity of the honey bee microbiome and the importance of comprehensive taxonomic representation for accurate classification (Liu et al., 2024; Marcelino et al., 2020; Wright et al., 2023), we used the NCBI Core Nucleotide Database (core_nt, 10/15/2025 version) as reference database. Although missing the intergenic regions, this database maximizes the taxonomic coverage against memory size, thus speeding up the classification process.

To maximize classification accuracy, we applied thresholds to the Kraken2 confidence score (0.25) and to the genus relative abundance, *i.e.* limit of detection (LoD, 5 × 10^-6^). These thresholds were determined using rarefaction (Schloss, 2024) and precision-sensitivity analyses, based on sets of known bee-associated taxa (Daisley & Reid, 2021), and effectively removed the effect of library size on taxon richness (Supp. Note 1).

The microbiome abundance table was constructed through the following steps: (i) Bracken output files were parsed and relative abundances values were computed (Supp. Note 1), (ii) relative abundances below the LoD were removed, (iii) only genera present in at least 5% of colonies were retained (Supp. Note 1) to reduce the number of statistical tests, focusing on the biologically most meaningful associations (Neu et al., 2021; Nickols et al., 2024), (iv) the filtered table was pivoted to have genera as columns and colonies as rows, and (v) missing or zero values, corresponding to abundances below the LoD, were imputed with ½ LoD (Mallick et al., 2021).

### *V. destructor*-related traits

For recapping and DMR, the transformed values with logit and empirical Bayes, respectively, from Eynard et al. (2024) were reused. For *V. destructor* load, in addition to the classical mite count phenotype (VL_count_) acquired by washing adult bees with detergent (Dietemann et al., 2013), we included a genomic-derived metric (VL_meta_) because it should be less affected by the observator and is readily available with such pooled sequencing data. We did not reuse the genomic-derived metric from Eynard et al. (2024) because it was not available for all colonies (Supp. Fig. 1A) and required an additional bioinformatics step. VL_meta_ consists in the log ratio between the number reads assigned to *Varroideae* and the number of honey bee reads, mapped to one of the three reference genomes or classified as *Apidae*. By relying on a log ratio transformation, VL_meta_ accounts for the compositional nature of metagenomics datasets (Gloor et al., 2017).

To satisfy the normality assumption of linear models, both *V. destructor* load measures were log-transformed after replacing zero values by 0.9 x the minimum non-zero value for each metric (Supp. Fig. 1A; Park, 2024). Indeed, zeros were interpreted as values below the LoD, *i.e.* sampling zeros, rather than true absence of mites, which is unlikely at colony scale. Moreover, this approach was found to maximize normality and reduce the effect of zeros on associations in preliminary tests.

### Microbiome models

To quantify the relative contribution of *V. destructor*-related traits (VL_count_, VL_meta_, recapping, and DMR) and environmental factors (season, year and location) to microbiome variation, we first visually inspected pairwise microbiome similarity heatmaps computed in R using the corr and pheatmap functions (R Core Team, 2026; Kolde R, 2025). Season was captured as a continuous variable representing the day of year (1–365) on which each colony was sampled, while location was defined by the apiary and the region (149 apiaries spread over 10 regions in France, and four other countries) where the colony was located.

We then searched for significant associations between these factors and the main axes of microbiome variations. These were identified by reducing the dimensionality of the microbiome abundance table using a principal component analysis (PCA). The associations between the main principal components (PC) and our factors of interest were modelled with linear mixed models implemented in the lme4 R package. To evaluate the contribution of *V. destructor*-related traits to microbiome variation, each trait was modelled individually with day of year, year, region as fixed effects, and apiary as a random effect such as in the following formula:

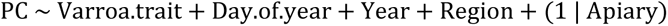

To correct for multiple testing, p-values were adjusted using the Bonferroni method by multiplying each p-value by the number of tested PC. Statistical significance across models was summarized and visualized using the pheatmap R package. To assess the influence of environmental factors on *V. destructor*-related traits themselves, these traits were predicted based on the environmental factors using the following formula:

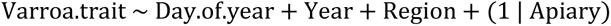

Next, we investigated which specific microbiome genera were associated with VL_count_, VL_meta_, recapping, and DMR, using linear mixed models implemented in MaAsLin 3 (Nickols et al., 2024). MaAsLin 3 tests associations at both the abundance and prevalence levels and partially accounts for compositionality by centering the slope (effect size) of each genus to the median slope across all genera. Given we were not particularly interested in the association between genera and regions, the following formula only included apiary to model the colony location:

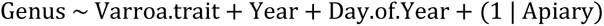

The microbiome abundance table and the transformed phenotypes and covariates were supplied as input feature and metadata tables, respectively, to the maaslin3 function. Because normalization and transformation had already been performed upstream, the normalization and transform parameters were set to NONE. The min_abundance and zero_threshold parameters were both set to log(LoD), while default values were retained for all other parameters.

## Results

### Large-scale metagenomic profiling of the European honey bee microbiome

To profile the European honey bee microbiome (Fig. 1), we conducted large-scale, reference-based metagenomics analyses of publicly available genomic data from 1,442 European honey bee colonies (Eynard et al., 2024). These colonies spanned 149 apiaries across 15 regions (10 regions in France, Switzerland, Netherlands, Luxembourg, Sweden) and were sampled once between June and October in 2016, 2017, or 2018. On average, 93.8% (SD: 2.4%) of the quality-filtered paired-end reads mapped to at least one of the three honey bee reference genomes (Supp. Fig. 3A). Of the remaining non-bee reads (mean: 2.2M per colony; SD: 1.0M), 61.3% (SD: 6.0%) were taxonomically classified (Supp. Fig. 3B; Lu et al., 2022).

*Bacteria* was by far the most abundant taxonomic group detected after honey bees, with a mean relative abundance (RA_mean_) of 0.034 (SD: 0.013; Fig. 3A), which accounted for 88.2% of the non-honey bee reads. *Bacteria* was also the most diverse group, with a mean genus richness of 25.4 (SD: 5.4) and representing an average of 74.1% (SD: 6.1%) of all genera detected per colony (Fig. 3B). The most abundant bacterial genera included four core gut microbiota members, *Lactobacillus*, *Gilliamella*, *Snodgrassella*, and *Bifidobacterium*, alongside genera typically described as non-core gut members: *Bartonella*, *Frischella*, *Commensalibacter*, and *Arsenophonus* (Fig. 3C). The five core genera, including *Bombilactobacillus* (RA_mean_: 1.4 × 10^-4^), as well as *Frischella* and *Commensalibacter* were detected in all colonies. *Bartonella* and *Arsenophonus* were found in 1413 and 891 colonies, respectively (Fig. 3C), and displayed the greatest inter-colony variability (Fig. 3D). The bacterial pathogenic genera *Spirosplasma*, which ranked among the ten most abundant bacterial genera, *Serratia*, and *Hafnia*, were observed in 739, 613, and 568 colonies, respectively. The etiological agents of European and American foolbroods, *Melissococcus plutonius* and *Paenibacillus larvae*, were detected in 141 and 22 colonies, respectively. *Ascosphaera apis*, causing Chalkbrood disease, was observed in 4 colonies.

**Figure 3.**
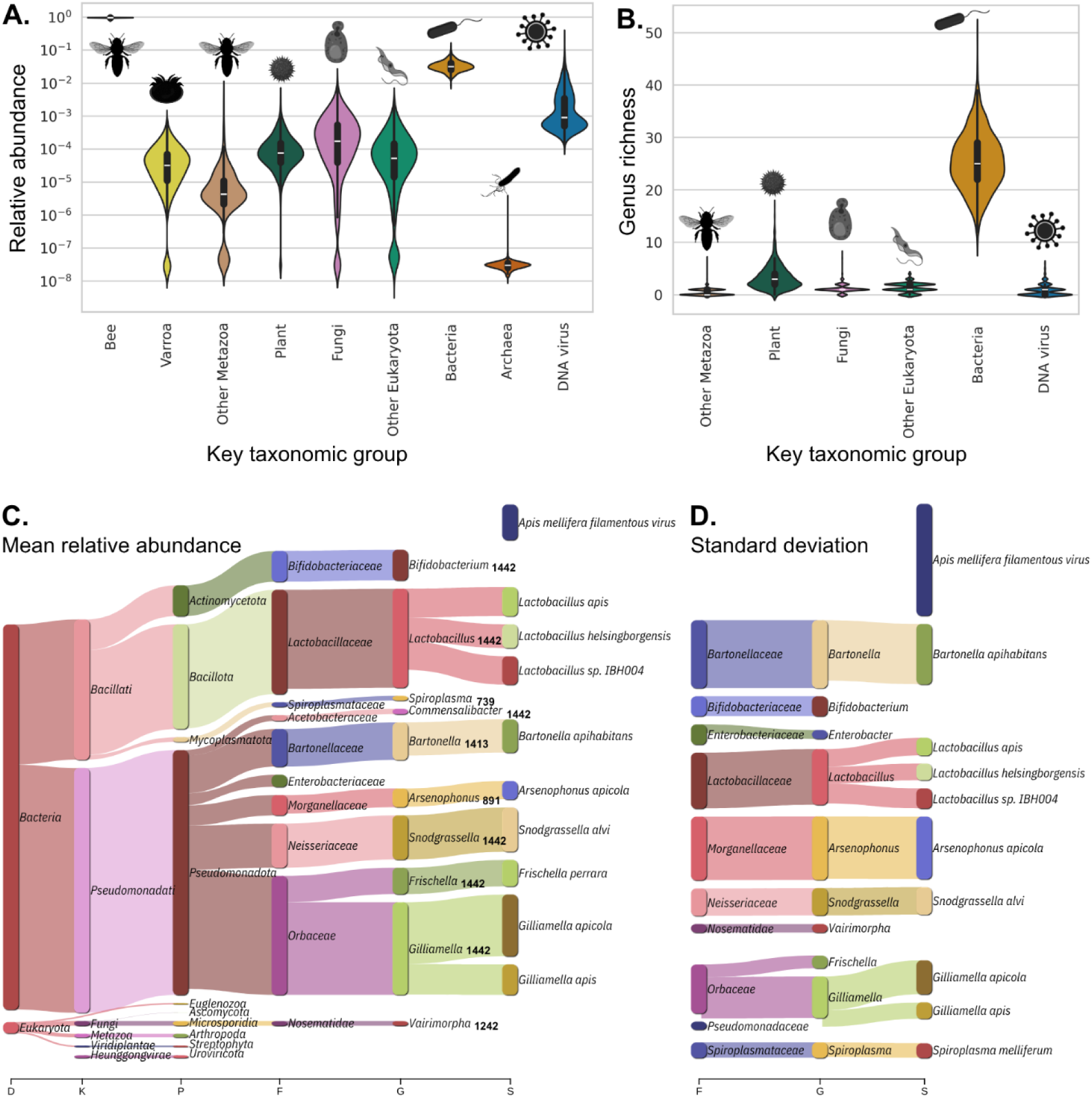
A) Relative abundance and B) diversity (genus richness) of key taxonomic groups, selected to represent the main components of the honey bee microbiome while covering all life, across 1,442 European honey bee colonies. Sankey plots generated using Pavian (Breitwieser & Salzberg, 2020), representing the top 10 (C) most abundant and (D) most variable taxa per taxonomic level. In C), bold figures show the number of colonies in which each genus was detected above the limit of detection.

DNA virus constituted the second most abundant non-honey bee taxonomic group (RA_mean_: 0.004; Fig. 3A), driven mostly by the *Apis mellifera* filamentous virus, which was also the most variable taxon across colonies (Fig. 3D). Bacteriophages, including *Caudoviricetes sp.*, represented the second most abundant viral group (Supp. Fig. 4). The remaining reads were eukaryotic (RA_mean_: 0.001) and consisted mainly of the known pathogens *Vairimorpha ceranae* (microsporidian; syn. *Nosema ceranae*) causing Nosemosis, *Lotmaria passim* (trypanosomatid), as well as residual honey bee sequences (Fig. 3C, Supp. Fig. 3). DNA from a wide range of flowering plants was also detected, as well as human DNA (Supp. Fig. 4).

Although relatively few reads mapped to *V. destructor* (Fig. 3A), they provided sufficient information to derive a new metagenomics-based metric for mite load (VL_meta_). VL_meta_ displayed a strong correlation with a previously described mtDNA-based mite load measure (Pearson *r* = 0.98) and with manual mite counts (Pearson *r* = 0.78, VL_counts_; Supp. Fig. 1B).

### Season, year, location, and *V. destructor* load contribute to microbiome variation but not resistance traits

We next assessed which factors (season, year, location, *V. destructor* load, recapping, and DMR) were associated with the variation in microbiome composition. To this end, we focused on the 80 genera detected above LoD (relative abundance of 5 × 10^-6^, Supp. Note 1) in at least 5% of colonies and combined visual inspection of pairwise microbiome similarity heatmaps with statistical modelling. Location and temporality emerged as the main factors associated with the microbiome of European honey bee colonies. Heatmaps, ordered geographically from south to north, revealed striking clustering patterns at both region and apiary levels (Fig. 4A). Some regions, such as Switzerland or Corsica (France) formed distinct microbiome clusters. Likewise, ordering colonies by sampling date revealed temporal structures (Supp. Fig. 5). These visual patterns were confirmed using linear mixed models fitted to the first six microbiome principal components (PC), which together explained over 50% of the total variance (Fig. 4B). Region (10 in France, and four other countries), sampling year, and season (sampling day) were significantly associated (q-value < 0.05) with five PC (Fig. 4C).

**Figure 4.**
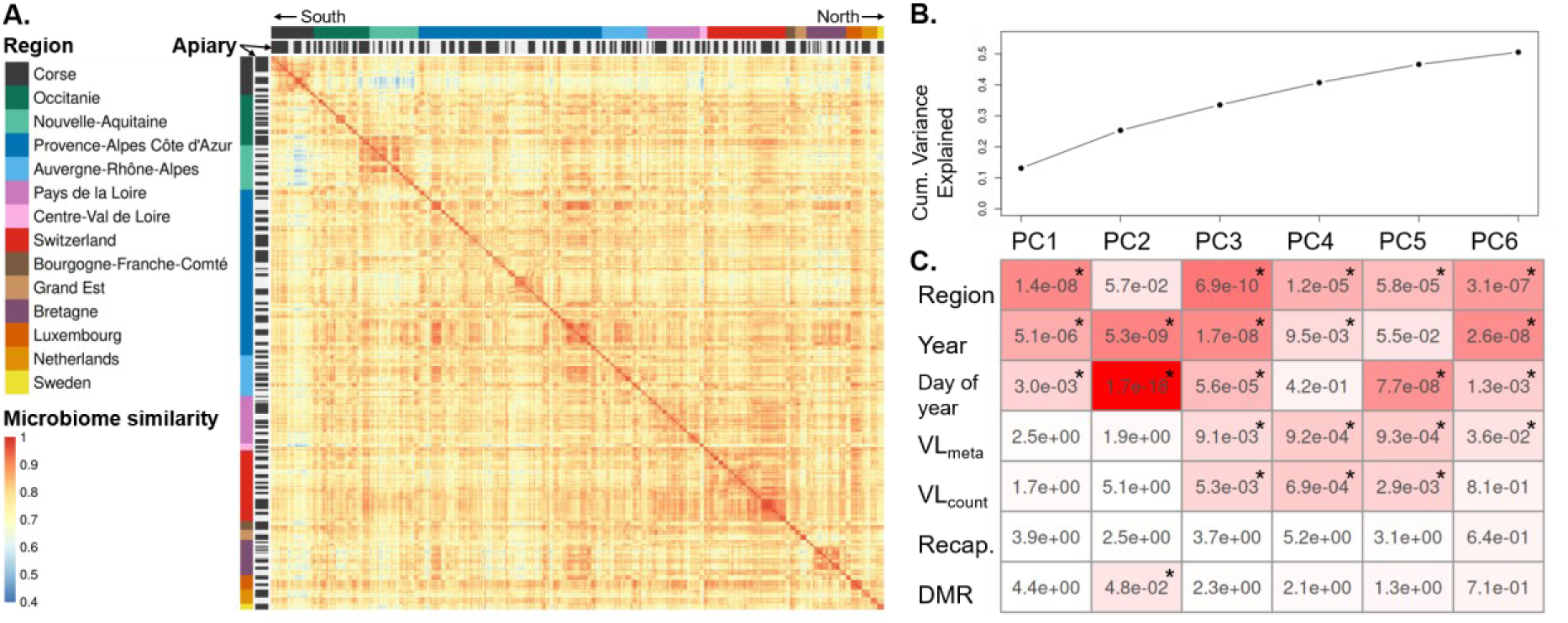
A) Heatmap displaying pairwise microbiome similarities between honey bee colonies. Colonies were ordered by apiary and geographically by region from south to north. Non-country regions are located in France. B) Cumulative variance explained by the first six principal components (PC). C) Significance of associations (q-values) between the microbiome-derived PC, covariates, and *V. destructor*-related traits. For categorical variables (region, year), only the lowest q-value is reported. Asterisks indicate significant q-values (<0.05).

*V. destructor* load, although to a lesser extent than location and temporality, was significantly associated with several microbiome PC (Fig. 4C). VL_meta_ was associated with four PC and VL_counts_ with three. By contrast, DMR was significantly associated only with one PC, and recapping showed no significant associations. No clustering was observed when correlation heatmaps were ordered by *V. destructor* load, recapping, or DMR (Supp. Fig. 6).

### Diverse microbiome genera correlate with *V. destructor* load

We then investigated the association between specific genera and *V. destructor*-related phenotypes using linear mixed models. Because season, year and location strongly influenced microbiome composition (Supp. Fig. 5, Fig 4) and correlated with *V. destructor*-related phenotypes (Supp. Fig. 2), these factors were included as covariates to avoid confounding associations.

Ten and 13 genera out of 80 were significantly associated (q-value < 0.05) with VL_meta_ and VL_count_, respectively (Fig. 5). Six genera were significantly associated with both *V. destructor* load metrics (highlighted with grey boxes in Fig. 5), and 5 genera identified significant with one metric were almost significant (q-value < 0.1) in the other one (highlighted with dashed grey boxes in Fig. 5, Supp. Fig. 7). Almost all significant genera, 9 for VL_meta_ and 11 for VL_counts_, were positively associated with *V. destructor* load, thus indicating increased abundance or occurrence probability at higher mite load. These genera included the pathogens *Morganella* (bacteria), *Lotmaria* (trypanosomatid), and *Vairimorpha* (microsporidian), as well as the non-core bacterial genera *Bartonella*, *Commensalibacter*, and *Entomomonas*. By contrast, only three genera were negatively associated with *V. destructor* load, and thus more prevalent or abundant in colonies with lower mite loads and belonged to the core gut bacteria *Bombilactobacillus*.

**Figure 5.**
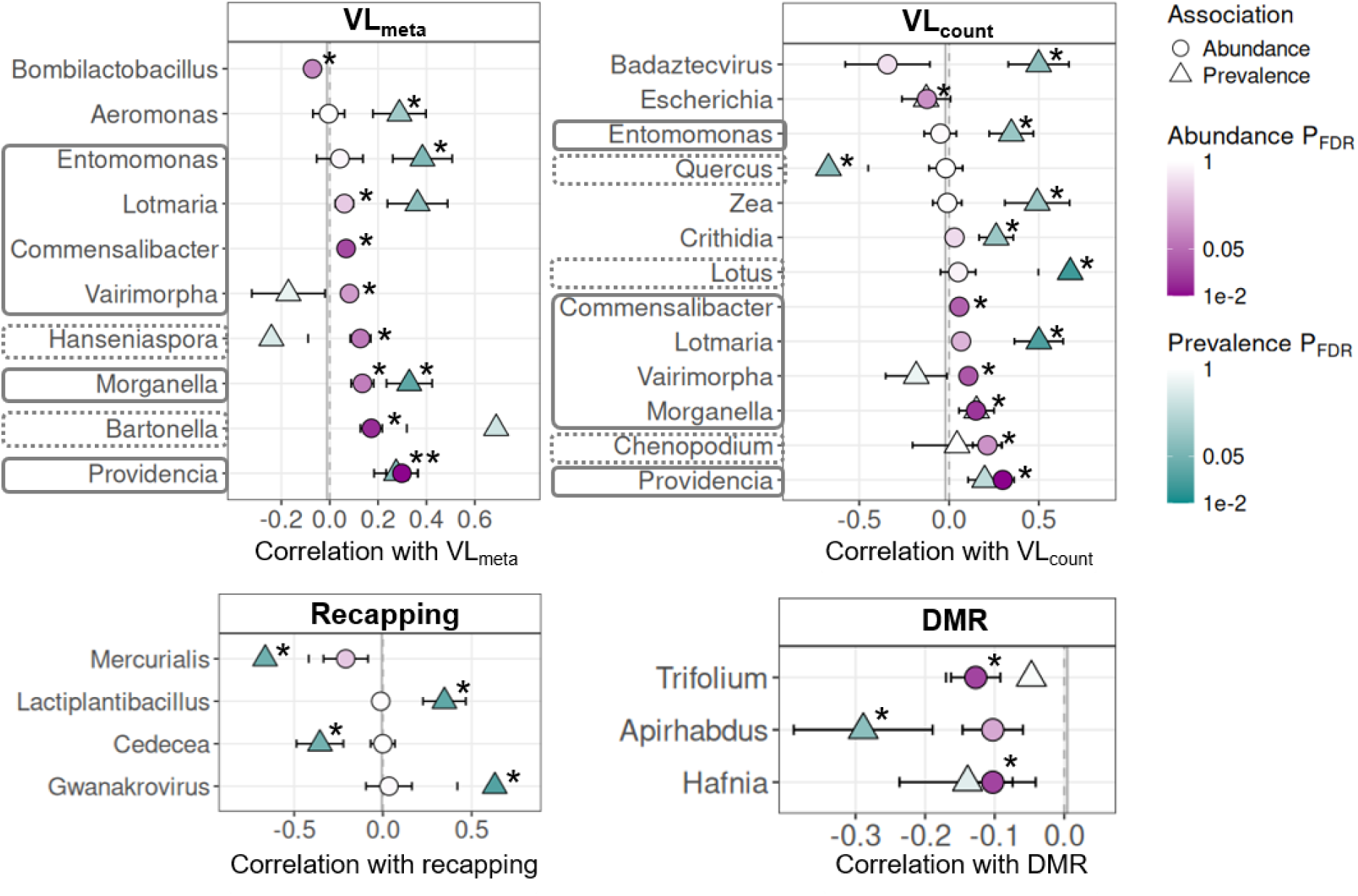
Microbiome genera significantly associated (q-value < 0.05, indicated by stars) with *V. destructor*-related phenotypes—*V. destructor* load (VL_meta_ and VL_counts_), recapping, and decreased mite reproduction (DMR)—inferred with linear mixed models. Grey boxes highlight genera associated with both *V. destructor* load metrics (both q-values < 0.05). Dashed grey boxes indicate genera significantly associated with one *V. destructor* load metric (q-value < 0.05) and marginally so (q-value < 0.1) with the other metric.

Consistent with the results of the PCA (Fig. 4C), recapping and DMR were associated with a substantially smaller fraction of the microbiome than *V. destructor* load. Specifically, only four genera were significantly associated with recapping and three with DMR (Fig. 5). The bacterial genus *Lactiplantibacillus* and the bacteriophage *Gwanakrovirus* (*Caudoviricetes*) showed positive correlations, while the genera *Cedecea*, *Apirhabdus*, *Hafnia* correlated negatively with these *V. destructor*-resistance traits.

Diverse plant genera also correlated with *V. destructor*-related phenotypes (Fig. 5). Quercus (oak trees), Lotus, Chenopodium, and *Zea* (incl. mays) were significantly associated with VL_count_, while the first three were close to the significance level using VL_meta_ (q-value < 0.1). *Mercurialis* and *Trifolium* were significantly associated with recapping and DMR, respectively. The associations identified in this section, which used log transformations of relative abundances, were highly consistent with two alternative compositionality-aware transformations (Supp. Note 2).

The abundance profiles of several genera associated with VL_meta_ were correlated (Supp. Fig. 8). Two non-core bacterial genera (*Entomonas*, *Bartonella*) and two pathogens (*Lotmaria*, *Vairimorpha*) formed the first cluster, while the second cluster comprised *Aeromonas* and two genera from the *Morganellaceae* family (*Providencia*, *Morganella*). Genera associated with VL_count_ formed less distinct clusters, consisting of the two trypanosomatids (*Lotmaria*, *Crithidia*) and the two *Morganellaceae* genera. The abundance profiles of the three genera associated with DMR correlated with each other, whereas those associated with recapping did not.

## Discussion

Although host-associated microbiomes play a central role in animal health, the general mechanisms driving microbiome variation and dysbiosis remain poorly understood, particularly in relation to parasitism and host resilience. Here, we present the largest metagenomics profiling of honey bee microbiomes, which we used to investigate the associations between parasitism by *V. destructor* and microbiome composition across 1,442 colonies. Using dimensionality reduction and statistical modelling, we show that *V. destructor* load, but not *V. destructor* resistance traits (recapping or DMR), is significantly associated with microbiome composition, although to a lesser extent than season, year, and location. By applying microbiome-tuned linear mixed models, we further identified numerous genera associated with *V. destructor* load, whereas only a few associated with resistance traits. While causal relationships cannot be established here, the scale of this dataset enables robust and generalisable conclusions about the interplay between *V. destructor* parasitism and honey bee microbiome composition.

By repurposing sequencing data originally generated for host genetic analyses, we demonstrate that meaningful microbiome profiles can be recovered even when more than 90% of reads map to host genomes. The remaining few million non-bee reads per colony were sufficient to recover a broad diversity of microbiome members, including the five core gut bacterial genera: *Bifidobacterium*, *Bombilactobacillus*, *Lactobacillus*, *Gilliamella*, and *Snodgrassella* (Motta & Moran, 2024). Although generally not described as core bacterial genera, *Frischella* and *Commensalibacter* were detected in all colonies, while the non-core genera *Bartonella*, and *Arsenophonus* were less frequent and displayed the greatest relative abundance variability. Bacterial pathogens were less prevalent. *Spirosplasma* spp., *Serratia* spp., and *Hafnia* spp. (Mouches et al. 1983; Raymann et al. 2018; Lang et al. 2022) were present in 40-50% of colonies, while the bacterial agents responsible for American and European foulbrood diseases, *P. larvae* and *M. plutonius* (Forsgren, 2010; Genersch, 2010), were found in 141 and 22 colonies, respectively. Our analysis also enabled us to identify non-bacterial pathogens including the microsporidian fungi *V. ceranae* causing Nosemosis (Higes et al., 2007) and the trypanosomid protists *Lotmaria passim* and *Crithidia mellificae*, that can damage gut epithelial tissues (Schaub, 1994). Surprisingly, *A*. *apis*, responsible for the relatively common chalkbrood disease (Aronstein & Murray, 2010), was observed in only four colonies, which could be due to the small set of sequences available in the reference database for this organism. The abundance of the DNA virus *Apis mellifera filamentous* virus was high and highly variable, while the presence of bacteriophages such as *Caudoviricetes* spp. could reflect their capacity to integrate into host bacterial genomes. In contrast to studies focusing on individual bee guts (Ellegaard & Engel, 2019; Prasad et al., 2025), comprehensive reference-based metagenomics analysis of pooled bee samples enabled us to capture the colony microbiome in a more holistic sense (Berg et al., 2020), encompassing the gut microbiota, microorganisms found in the honey bee haemolymph, environmental DNA found on the surface of honey bees or in their digestive tract such as pollen, as well as *V. destructor* mites and their microbiome (Fig. 1).

Another methodological product of this approach is the metagenomics-based *V. destructor* load phenotype VL_meta_, derived from the ratio of reads assigned to *V. destructor* relative to honey bee reads. Despite its simplicity, VL_meta_ showed strong agreement with manually measured mite counts (Pearson *r* = 0.78) and an almost complete concordance with a previously established metric based on the ratio between *V. destructor* mtDNA and honey bee genomic DNA (Pearson *r* = 0.98; Eynard et al., 2024). These results suggest that metagenomics-derived mite load estimates could provide a scalable, standardised, and observer-independent alternative phenotype to traditional count-based methods. Future work should evaluate the repeatability and potential heritability of VL_meta_, particularly in the context of breeding programs for *V. destructor* resistance (Guichard et al., 2020).

With 1,442 colonies spanning five European countries, five months and three years—30 times more than in the largest study to date by Hubert et al. (2017)—this dataset provides a statistically robust and generalisable assessment of the contribution of temporal, spatial, and parasitic drivers to honey bee microbiome variation. Consistent with previous work (Hroncova et al., 2015; Kešnerová et al., 2020; C. Li et al., 2022), location and temporality emerged as important factors shaping microbiome composition. Indeed, sampling season, year, and location were significantly associated with five out of the six microbiome-derived PC that explained more than 50% of the microbiome variance. *V. destructor* load showed a more modest but significant association with the microbiome, correlating with four microbiome-derived PC and with approximately 20% of genera. We believe this result is particularly robust given it was reached with two metrics of *V. destructor* load (VL_meta_ and VL_count_, Fig. 4, Fig. 5), and that three different compositionality-aware transformations of genus relative abundances yielded highly consistent genus-level associations (Supp. Note 2). While confirming that mite infestation is not the primary factor influencing microbiome variation in honey bee workers (Y. Wang et al., 2025), our results, thanks to the large size of our dataset and proper covariate modelling, demonstrate the link between *V. destructor* parasitism and honey bee microbiome composition.

The identity of the genera associated with *V. destructor* loads and the direction of their correlation with mite infestation levels provide ecological insights into host-microbiome interactions in honey bees. The majority of significant genera correlated positively with *V. destructor* load and comprised known pathogens and potential opportunists, whose co-varying abundances suggest that *V. destructor* infestation is linked not to isolated taxa but to broader pathogenic assemblages. Among these, the trypanosomatids (*Lotmaria* and *Crithidia*) and the microsporidian *Vairimorpha* are established honey bee pathogens. The bacterial genus *Morganella* includes *Morganella morganii*, recently shown to be vectored by *V. destructor* and highly lethal to honey bee colonies (Chen & Huang, 2025), while the related genus *Providencia* (family *Morganellaceae*), though not comprising described honey bee pathogens, contains insect pathogen species (Galac & Lazzaro, 2011). Several non-core potentially opportunistic taxa also correlated positively with mite load. While associations with *Bartonella* and *Commensalibacter* have been previously reported (Hubert et al., 2017; Y. Wang et al., 2025), this study also provides evidence of a link with the yeast *Hanseniaspora* (Agarbati et al., 2024) and the bacteria *Entomomonas* (family: *Pseudomonadaceae*), a genus recently described in *Apis cerana* whose reduced genome—particularly regarding carbohydrate metabolism—suggests an intimate relationship with honey bees (J. Wang et al., 2020). By contrast, *V. destructor* load correlated negatively with the core bacteria *Bombilactobacillus* (family *Lactobacillaceae*), such that higher mite infestation levels were associated with reduced abundance of this core symbiont. The beneficial impact of *Lactobacillaceae* spp. for mite control has already been experimentally demonstrated (Pitek et al., 2025; Tejerina et al., 2020). Taken together, the positive correlations between mite load and pathogens or opportunists, alongside the negative correlation with the core symbiont *Bombilactobacillus*, suggest that parasitism by *V. destructor* is associated with a shift in the composition of the honey bee microbiome toward dysbiosis (Maes et al., 2016). Although this pattern may partly reflect the increased abundance of the mite’s own microbiome in colonies with higher mite loads, it cannot explain the decline in the core symbiont, *Bombilactobacillus*. These findings are compatible with several non-exclusive explanations. Colonies harbouring unbalanced microbial communities may be inherently more vulnerable or attractive to mite infestation. Conversely, *V. destructor* infestation may itself disrupt the microbiome, either directly through microbial transfer during feeding, or indirectly by weakening the colony (Pusceddu et al., 2025; Warner et al., 2024) and thereby increasing its susceptibility to pathogenic and opportunistic taxa (Hubert et al., 2017).

In contrast to *V. destructor* load, we found limited evidence for associations between microbiome composition and *V. destructor* resistance traits. Nonetheless, one microbiome-derived PC was significantly associated with DMR and *Lactiplantibacillus* (family: *Lactobacillaceae*) correlated positively with recapping. Interestingly, *Lactiplantibacillus* has been linked to behavioural changes in mice, while *Lactobacillaceae* to olfactory memorisation and learning in honey bees (W.-H. Liu et al., 2016; Zhang, Mu, Cao, et al., 2022)—sensory capacities that are likely essential for the detection of *V. destructor*-infested brood cells (Mondet et al., 2020; H. Wang et al., 2026). Moreover, hygienic-performing bees have been shown to harbour higher abundances of *Apilactobacillus kunkeei*, another *Lactobacillaceae* member, and this family is involved in immune modulation (Motta & Moran, 2024; Tola et al., 2025). Individuals performing resistance behaviours may therefore benefit from increased *Lactobacillaceae* abundance both for pathogen defence and enhanced sensory detection. While the scarcity of observed associations between the honey bee microbiome and *V. destructore* resistance traits is plausible, methodological limitations may also explain these results. Short-read metagenomics, combined with a relatively low proportion of microbial reads, constrained our ability to resolve finer-grained taxonomic variation, particularly at species and strain levels. This variation is known to be functionally important within core gut lineages (Ellegaard & Engel, 2019). The pooling of workers also masks individual-level microbiome variation, which may be more informative for detecting associations with behavioural traits (Tola et al., 2025). Furthermore, the absence of RNA sequencing data precluded the detection of RNA viruses such as DWV, which is a major cause of colony decline closely associated with *V. destructor* infestation (Beaurepaire et al., 2020; Martin & Brettell, 2019). Future studies would therefore benefit from microbial DNA enrichment, metatranscriptomics, long-read sequencing, and assembly-based metagenomic methods to achieve greater taxonomic resolution.

The association between multiple plant genera and *V. destructor* load, recapping, and DMR suggests that the availability of floral resources may play a role in the colonies varying ability to resist *V. destructor*. Pollen and nectar contain diverse secondary metabolites that pollinators can use to resist parasites (Manson et al., 2010) and reduce virus loads (Palmer-Young et al., 2017). Beyond these direct pharmacological effects, the abundance and quality of food itself (*i.e.* pollen and nectar availability and composition) has been linked to improved colony survival, reduced *V. destructor* load and immunity (Requier et al., 2017). This suggests that differential access to floral resources across plant communities could underlie the observed associations. Diet is also a known driver of honey bee microbiome composition (Hroncova et al., 2015; Kešnerová et al., 2020; C. Li et al., 2022; Meehan & O’Toole, 2025), raising the possibility that plant-mediated shifts in microbiome composition indirectly influence colony resistance to *V. destructor* via the microbiome role in honey bee health (Raymann & Moran, 2018). However, the low quantity of plant-derived DNA in our dataset limited our ability to characterise floral diversity at fine taxonomic resolution, and this question should be further explored in future studies.

In conclusion, while this study does not attempt to establish causality, it is the largest metagenomic analysis of honey bee microbiomes to date and provides robust, generalisable evidence of associations between microbiome composition, season, year, location, and *V. destructor* parasitism. The identified microbial taxa constitute valuable candidates for experimental work aimed at establishing causal mechanisms and developing microbiome-based strategies to enhance honey bee colony resilience. Beyond ecological insight, these findings carry practical implications: microbiome composition may serve as a marker of colony health and as a complementary criterion in selective breeding programs targeting *V. destructor* resistance, ultimately contributing to more sustainable honey bee health management.

## Acknowledgements

We would like to thank Sonia Eynard, Benjamin Basso, and Alain Vignal for providing valuable additional information regarding the samples and the broader context of the BeeStrong project, as well as for fruitful discussions. We are also grateful to Julia Gustavsen, Julien Brunisholz, and the entire Scientific IT team of Agroscope for providing computational resources and support. Finally, we thank Charlotte Huyghe for her advice on metagenomics analyses and Tanja Herrmann for her valuable input in designing the figures.

## Author contributions

VR, BD, VD, MN, and PE conceptualized the study; VR analysed the data and create the figures; TL assisted with data analysis; BD, VD, and MN acquired funding for the study; VR wrote the first draft; all authors read, edited, and agreed to the final version; BD managed the project.

## Conflict of interest

The authors declare that they have no conflict of interest.

## Funding

This work was supported by the Swiss National Science Foundation (SNSF) [Grant Number 10000808].

## Data Availability

Raw sequences were made available by Sonia Eynard and collaborators (2024) under the bioproject accession PRJNA1083455. The original metadata analysed in this article is available here: https://doi.org/10.57745/HHE4CZ. Scripts and cleaned metadata used in this study are deposited to Zenodo: https://doi.org/10.5281/zenodo.21195665.

## Notes

https://doi.org/10.5281/zenodo.21195665

https://doi.org/10.57745/HHE4CZ

### Competing Interest Statement

The authors have declared no competing interest.

