## Supplementary material for "Temporal, spatial, and parasitic drivers of microbial variation in European honey bees"

#### Supplementary figures

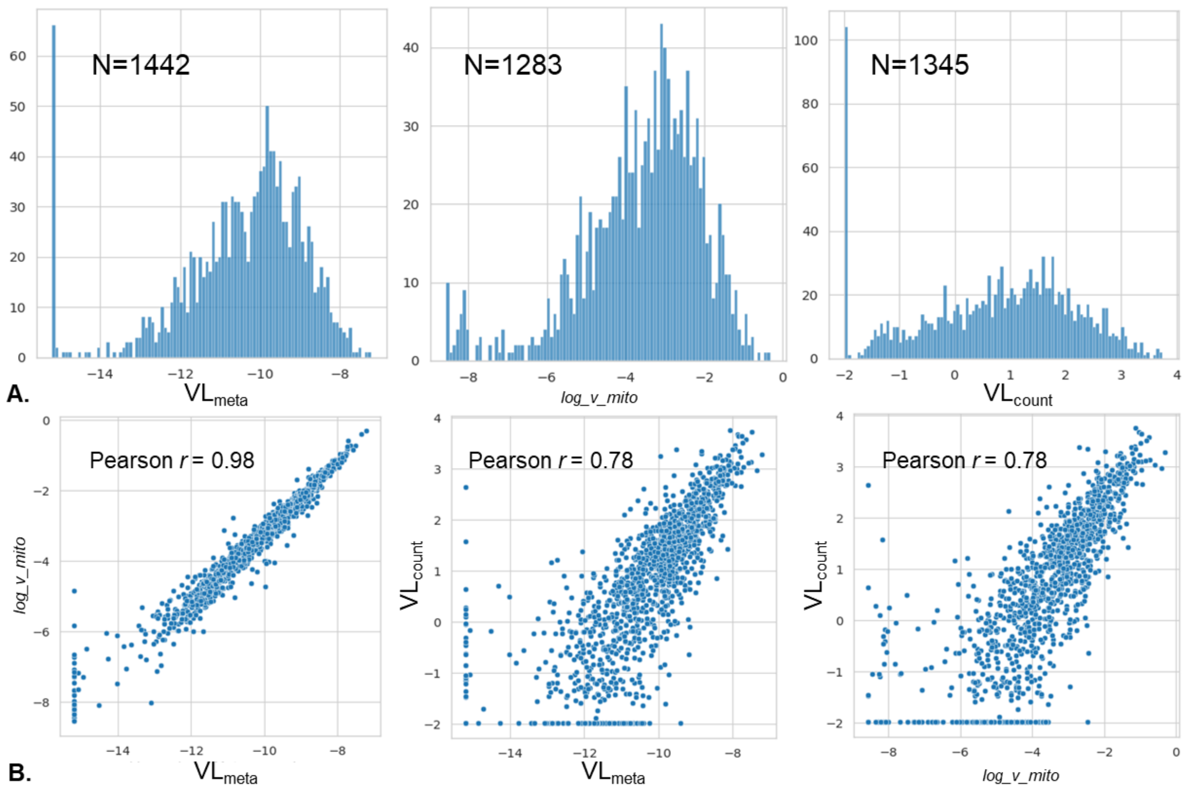


**Supp. Figure 1.** A) Distribution of *V. destructor* load phenotypes and number of colonies with a value. B) Correlations between *V. destructor* load phenotypes. $VL_{count}$ *V. destructor* load quantified by washing adult bees with detergent $VL_{meta}$ genomic-derived metric of *V. destructor* load; log_v_mito: genomic-derived metric from Eynard et al. (2024).

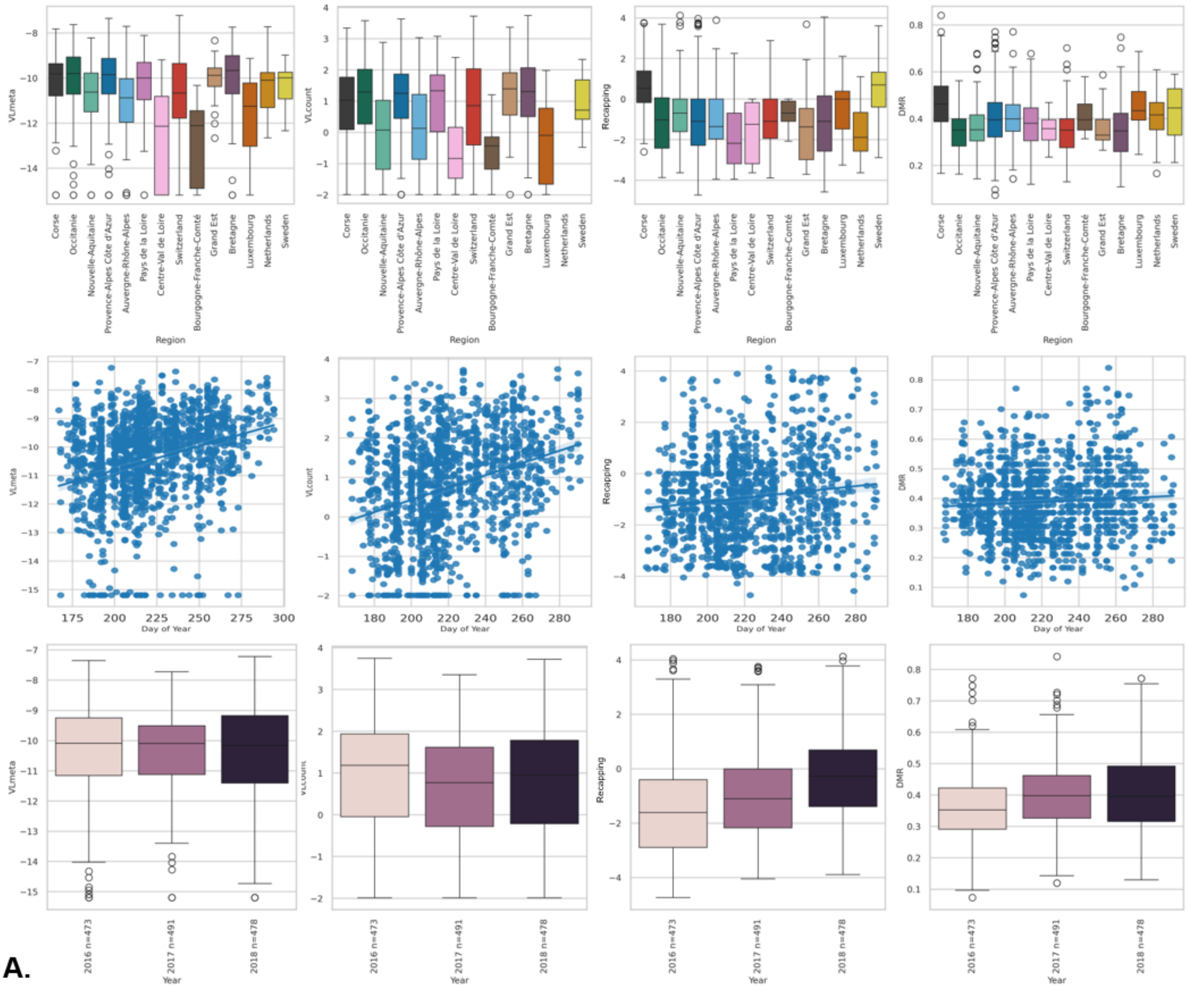


**A.**

| | Region | Year | Day of year |
|---|---|---|---|
| $VL_{meta}$ | 4.1e-02* | 3.5e-02* | 6.9e-12* |
| $VL_{count}$ | 8.5e-02 | 7.8e-01 | 7.3e-12* |
| Recap. | 2.3e-08* | 4.0e-12* | 9.8e-01 |
| DMR | 7.8e-10* | 4.0e-11* | 2.4e-01 |

**B.**

**Supp. Figure 2.** A) Correlations between *V. destructor*-related traits and environmental covariates. B) Associated p-values. Recap: Recapping; DMR: Decreased Mite Reproduction; $VL_{count}$: *V. destructor* load quantified by washing adult bees with detergent; $VL_{meta}$: genomic-derived metric of *V. destructor* load.

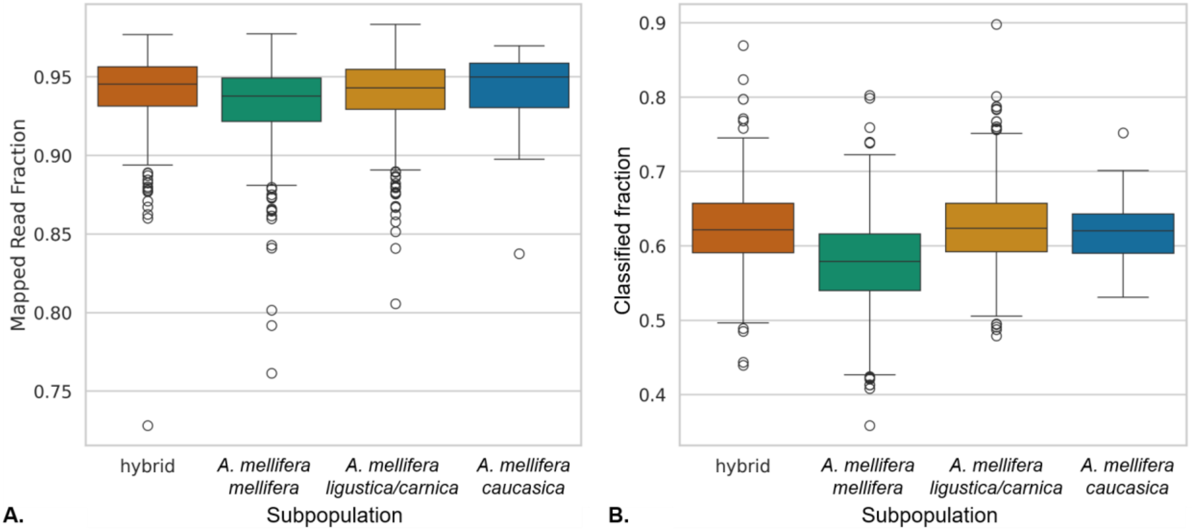


**Supp. Figure 3.** A) Mapping rate to at least one of the three honey bee reference genomes using Bowtie2, B) Proportion of non-bee reads classified with Kraken2 and Bracken.

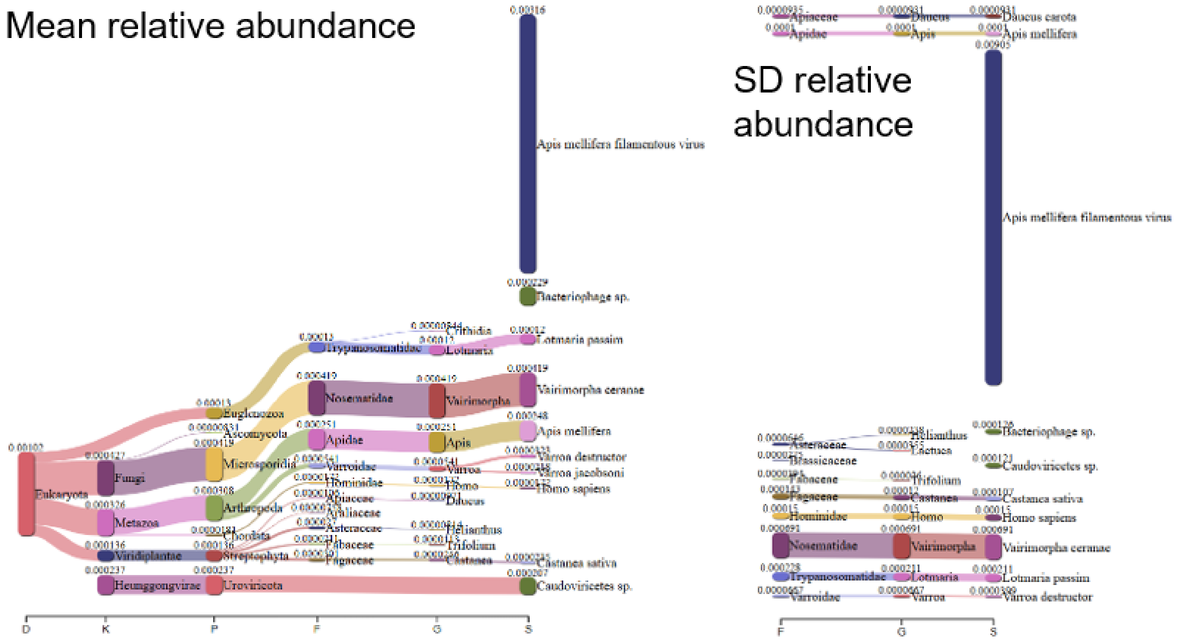


**Supp. Figure 4.** Sankey plots generated using Pavian (Breitwieser & Salzberg, 2020) representing the top 10 non-bacterial most abundant (left, mean relative abundance) and variable (right, SD relative abundance) taxa per taxonomic level.

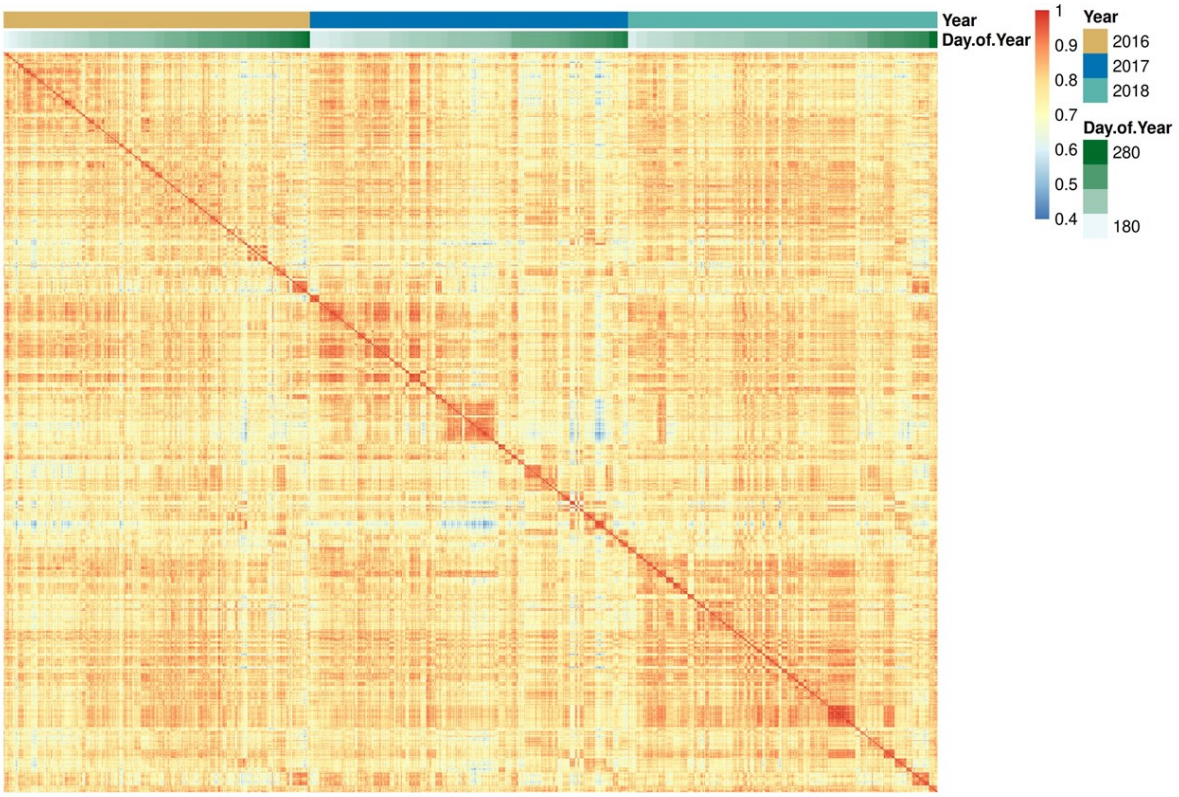


**Supp. Figure 5.** Heatmap displaying pairwise microbiome similarities between honey bee colonies ordered by sampling year and day of year.

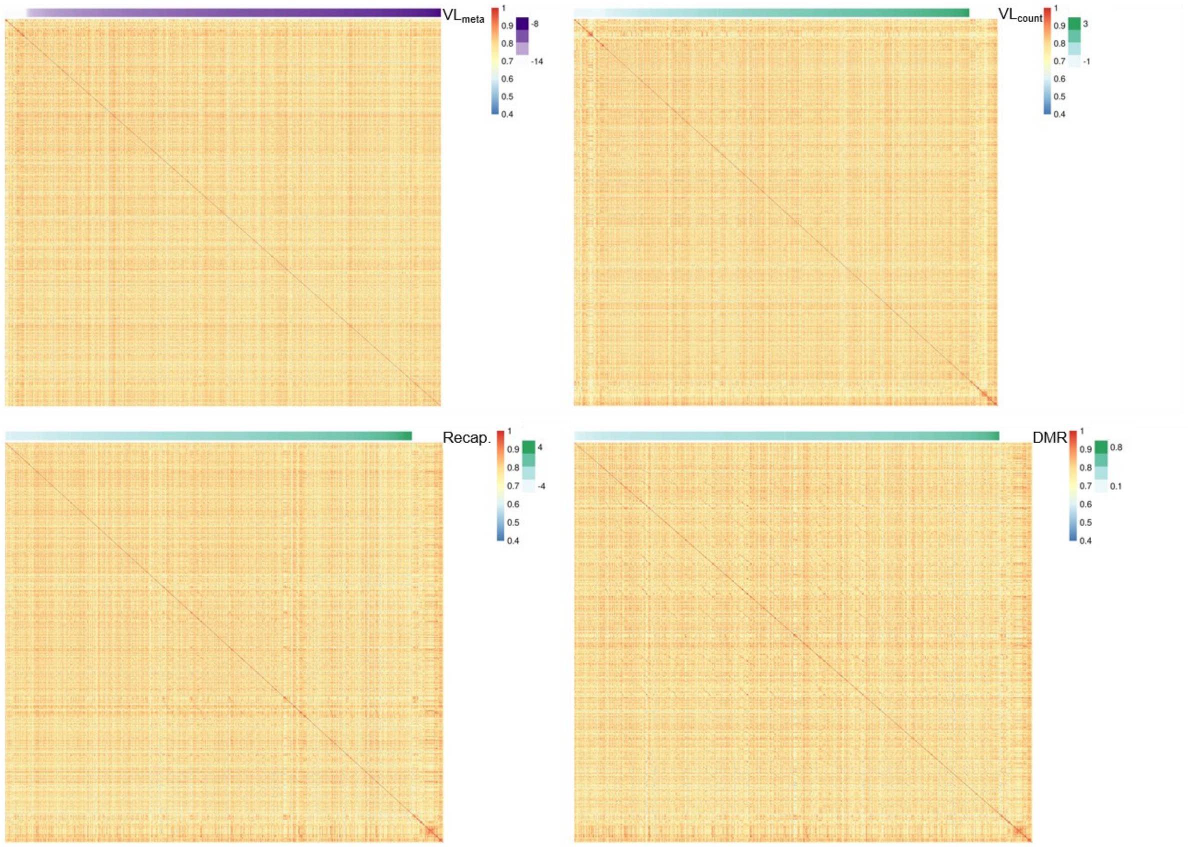

**Supp. Figure 6.** Heatmaps displaying pairwise microbiome similarities between honey bee colonies ordered by *V. destructor*-related trait value. Recap: Recapping; DMR: Decreased Mite Reproduction; $VL_{count}$: *V. destructor* load quantified by washing adult bees with detergent; $VL_{meta}$: genomic-derived metric of *V. destructor* load.

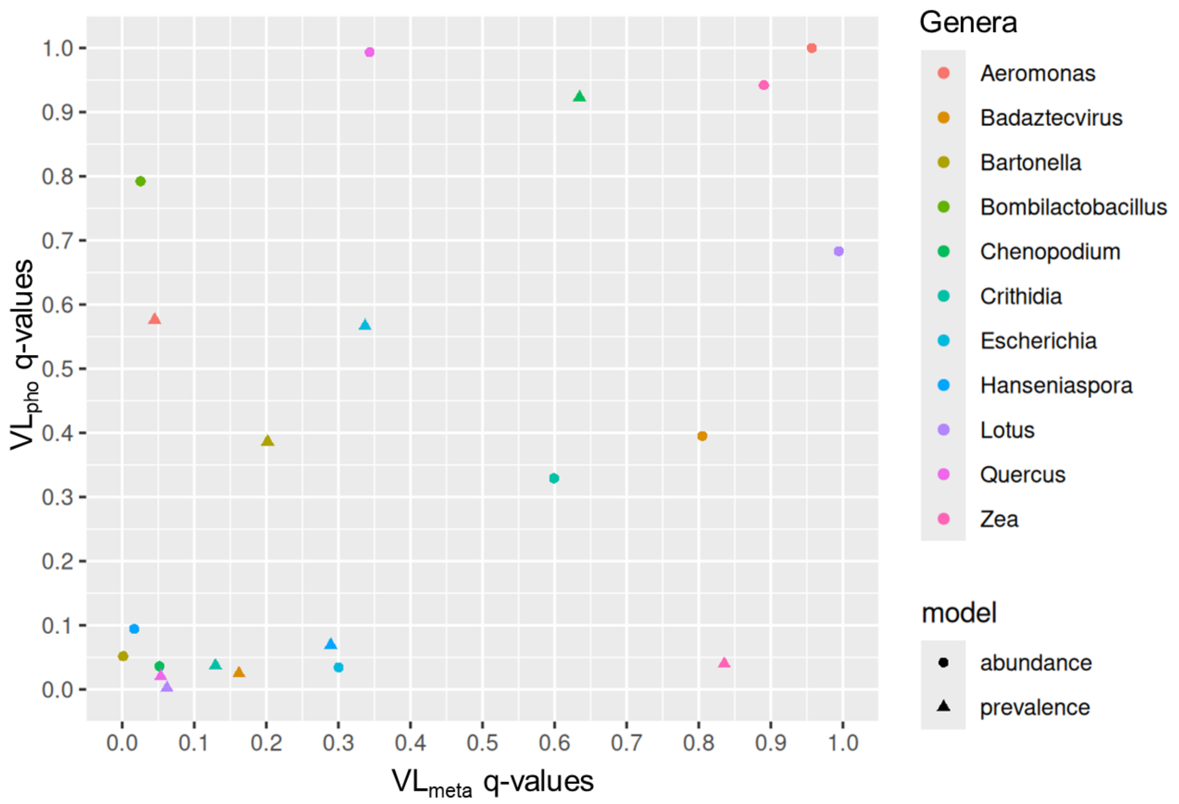


**Supp. Figure 7.** Comparison between q-values obtained with the two *V. destructor* load. $VL_{count}$: *V. destructor* load quantified by washing adult bees with detergent; $VL_{meta}$: genomic-derived metric of *V. destructor* load. Only genera found significantly associated with only one metric are displayed.

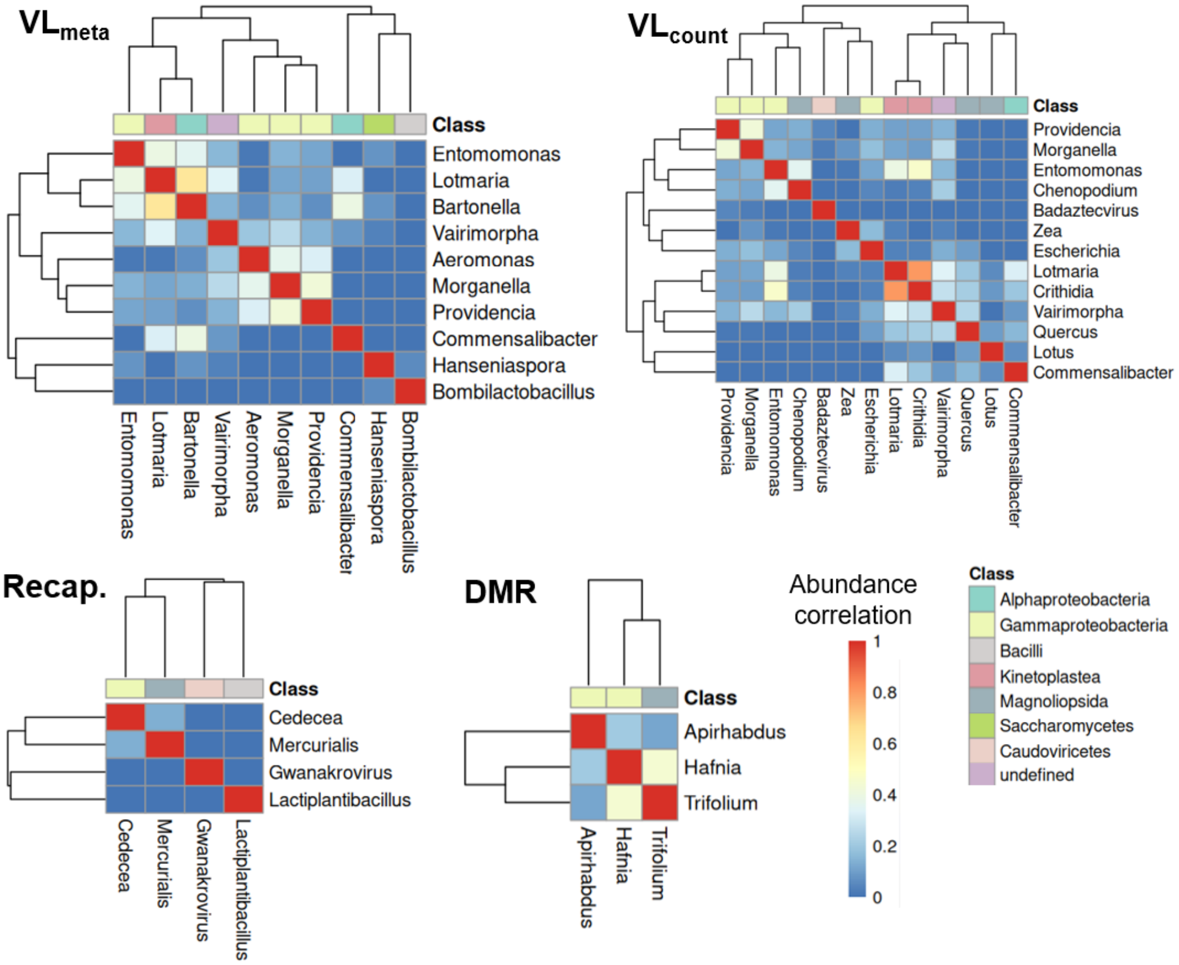


**Supp. Figure 8.** Heatmaps of pairwise similarities between abundance profiles of microbial genera significantly associated with *V. destructor* load, recapping, and decreased mite reproduction (DMR). Recap: Recapping; DMR: Decreased Mite Reproduction; $VL_{count}$: *V. destructor* load quantified by washing adult bees with detergent; $VL_{meta}$: genomic-derived metric of *V. destructor* load.

### Supplementary Note 1: Calculating relative abundances and finding the appropriate thresholds for taxonomic profiling

To control for the effect of library size on genus abundance, relative abundances were calculated by dividing Bracken-estimated read counts by the total number of reads with known taxonomic identity, thus either aligned to at least one bee genome or classified by Kraken2 and Bracken. Including host reads in the denominator rather than merely microbial reads was motivated by the high and relatively constant abundance of honey bee DNA across samples. This approach may help mitigate the effects of the compositional nature of metagenomic data because the number of honey bee cells, and thus DNA molecules, should be relatively constant across samples (Gloor et al., 2017; Tsilimigras & Fodor,

2016). Reads unclassified by Kraken2 or Bracken were excluded from the denominator to ensure that relative abundances summed to one within each sample. Note that bee relative abundances were calculated by combining the proportion of reads classified as bee, clade *Anthophila*, with the ones mapping to one of the three honey bee reference genomes.

Because relative abundances alone do not correct for library size biases on diversity estimates such as genus richness and do not control for false positive taxonomic assignments (Weiss et al., 2017), we applied thresholds on both the Kraken2 Confidence Score (CS) and genus relative abundances—heron Limit of Detection (LoD). While the former minimizes false positive read assignments by ensuring that a minimum fraction of read *k*-mers map uniquely to the assigned genus, the LoD choice directly impacts the false positive rate of the taxonomic classification by ensuring that enough reads support the detection of a genus. These thresholds were identified using rarefaction analyses and precision-sensitivity analysis based on sets of known bee associates.

Genus rarefaction curves were computed on a subset of 20 colonies. Kraken2 output files were subsampled ten times at different subsampling depths and Bracken was rerun for each subsample. For subsamples to have the same library size (*i.e.* number of reads) across colonies, the library size of the largest subsample was constrained to the one of the smallest colony. Thresholds combinations yielding rarefaction curves that plateaued near the largest subsample size were retained as candidates. Indeed, a premature plateau indicates that true taxa are probably missed, whereas non-converging rarefaction curves suggest residual library size bias. As a complementary analysis to assess precision and sensitivity of the different candidate CS-LoD threshold pairs, we used a set of taxa known to be associated with *Apoidea* (i.e. not specific to honey bees) from the BEExact database v2023.01.30 (Daisley & Reid, 2021). Considering the numbers of true positives (TP), false positives (FP) and false negatives (FN), precision was defined as the proportion of detected taxa that are known bee associates (TP / [TP + FP]), and sensitivity as the proportion of known bee-associated taxa recovered within a sample (TP / [TP + FN]). F1 is the harmonic mean between precision and sensitivity.

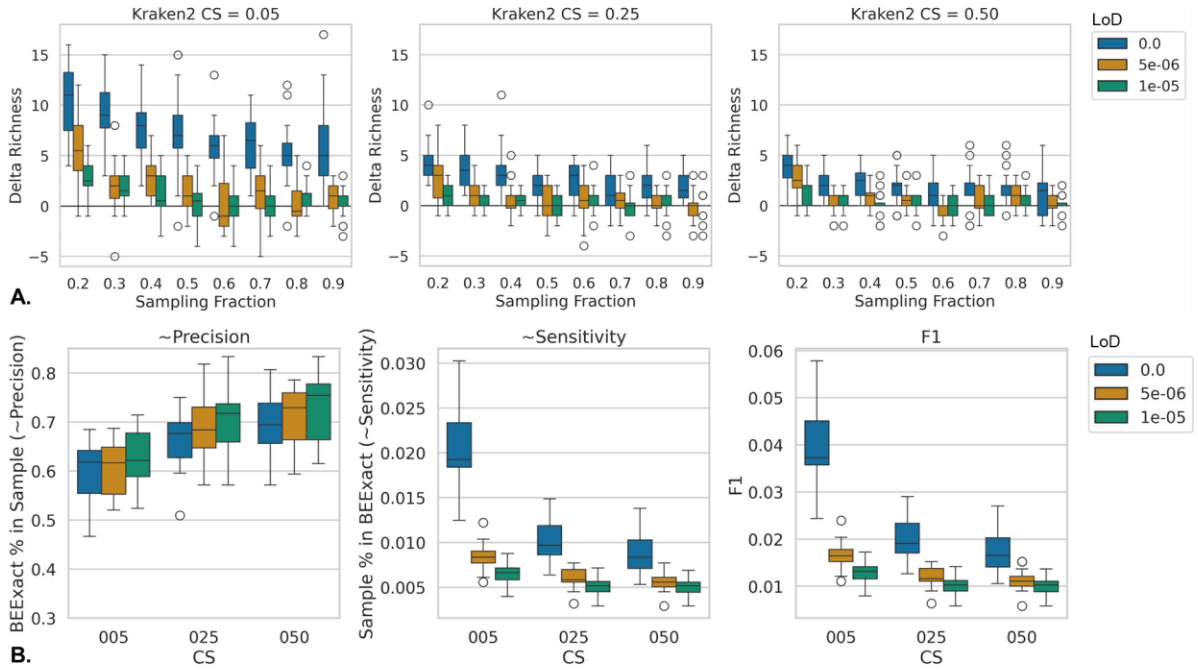


**Figure 1.** A) Aggregated rarefaction curves for 20 colonies. Delta richness indicates the number of new genera detected compared to the previous subsampling fraction. B) Precision-sensitivity analyses, based on sets of known *Apoidea*-associated taxa. CS: Kraken2 confidence score; LoD: Limit of Detection; F1: harmonic mean between precision and sensitivity.

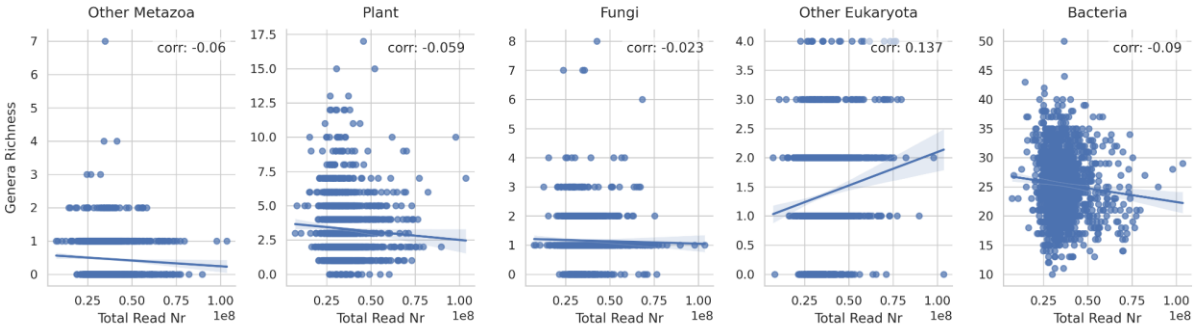


**Figure 2.** Correlations between library sizes (Total Read Nr) and genera richness at the LoD and CS thresholds used in this study.

These analyses indicated that increasing the Kraken2 CS to 0.25 strongly increases the proportion of bee-associated taxa identified in colonies (~precision, Fig. 1B). Moreover, a modest LoD threshold of $5 \times 10^{-6}$ relative abundance was found to reduce library size effects on genus richness (Fig. 1A). Further increases of CS or LoD were not found to have significant effects on rarefaction convergence nor on classification precision (Fig. 1). Similar modest CS values have been recommended in benchmarking studies to limit the number of unclassified reads and maximize the congruence between observed and true relative abundances (Y. Liu et al., 2024; Wright et al., 2023). The choice of these thresholds was validated across all colonies by the limited residual correlation remaining between taxon richness and sequencing depth across all domains of life (Fig. 2).

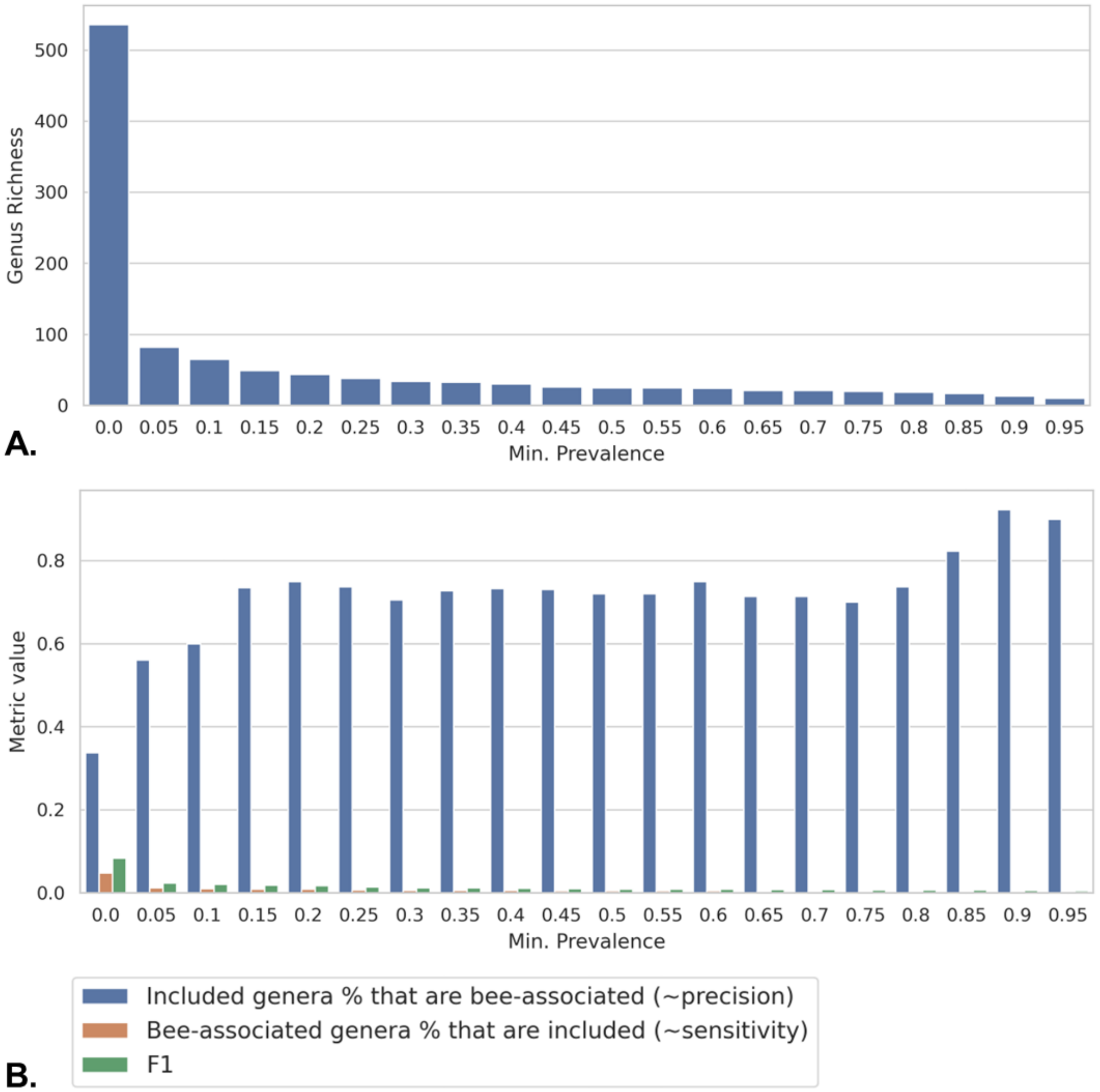


**Figure 3.** A) Genus richness at diverse prevalence cutoff values. B) Precision, sensitivity, and accuracy (F1) values at different prevalence cutoff values.

To determine an appropriate prevalence threshold, we examined how genus richness and the proportion of bee-associated genera changed across a range of cut-off values (Fig. 3). Restricting analyses to genera present in at least 5% of colonies was supported by two observations: genus richness declined steeply with increasing threshold values, and the proportion of bee-associated genera was substantially higher than at lower cut-off values. Applying this filter also had the practical benefit of reducing the number of statistical tests.

### Supplementary Note 2: Comparing Log Relative Abundances to compositionality-aware transformations

MaAsLin 3 partly accounts for compositionality after the Log relative abundance (LRA) transformation. Nonetheless, we repeated all analyses with two common transformations used to correct for compositionality: the Additive Log-Ratio (ALR), using honey bee as the reference, and the Centered Log-Ratio (CLR), which normalizes taxon abundances by their geometric mean (Gloor et al., 2017). For ALR, the denominator was the total number of honey bee reads—mapped to one of the three reference genomes or classified as *Apidae*. To ensure comparable limits of detection (LoD) across transformations, we fitted linear regressions between LRA and the corresponding CLR or ALR values, using the predicted value at log($5 \times 10^{-6}$) as the LoD for each (ALR: −12.17, CLR: 4.73). Results were highly consistent across all three transformations (Fig. 1).

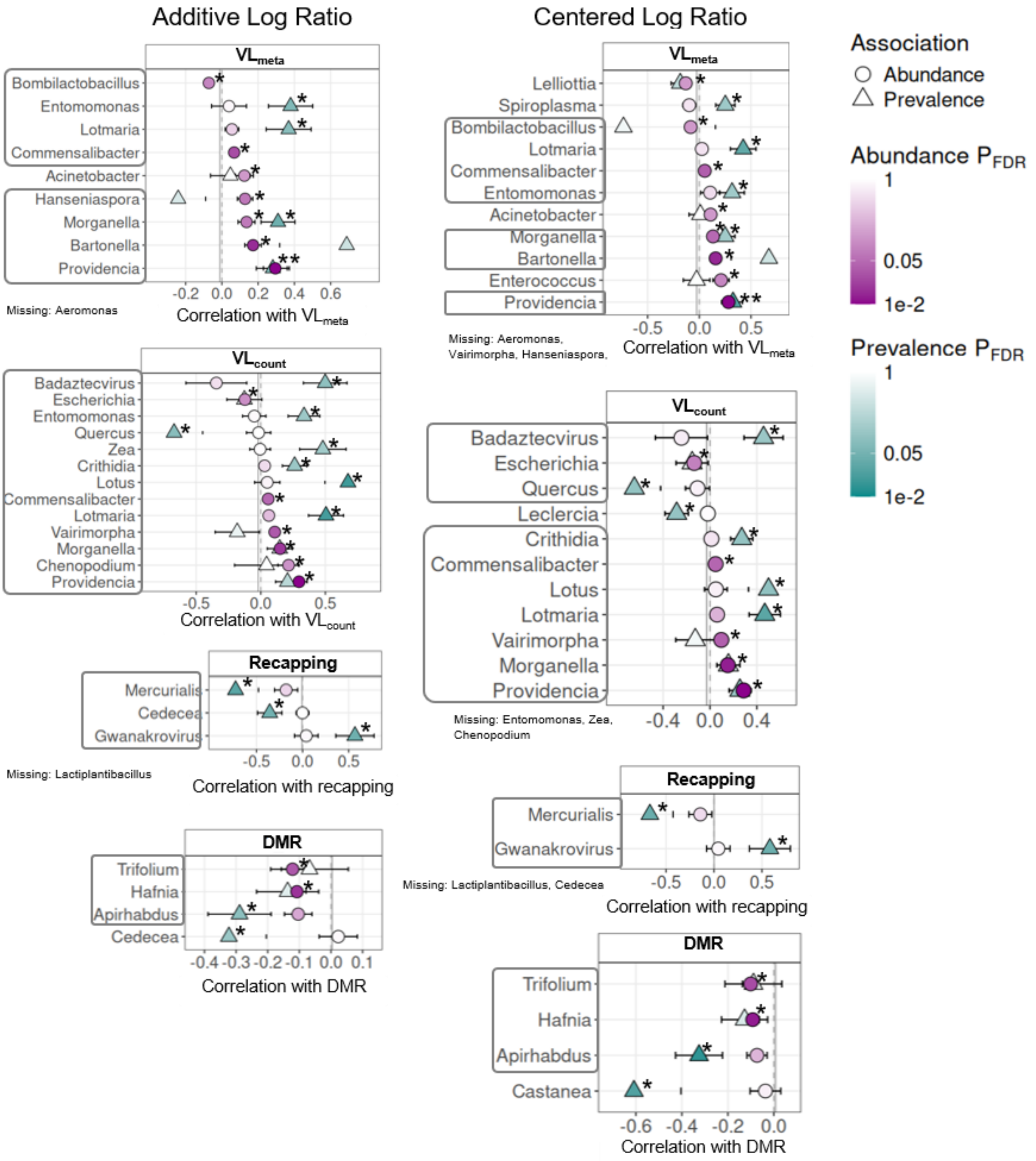

**Figure 1.** Microbiome genera significantly associated (q-value < 0.05, indicated by asterisks) with *V. destructor*-related phenotypes—*V. destructor* load ($VL_{meta}$ and $VL_{counts}$), recapping, and decreased mite reproduction (DMR)—inferred with linear mixed models. (Left) Additive Log Ratio transformation. (Right) Centered Log Ratio transformation. Grey boxes highlight significant genera found with Log Relative Abundance transformation (Main text, Fig. 5).
